# The gender gap in academic career achievements and the mediation effect of work-family conflict and partner support

**DOI:** 10.1101/2022.03.23.485507

**Authors:** Xiang Zheng, Haimiao Yuan, Chaoqun Ni

## Abstract

Parenthood has long been associated with gender disparities in academia. Yet, the underlying mechanism of how parenting is associated with career achievement gaps of academics remains unclear. Using data from a large-scale survey distributed to 7,764 scholars and their publication profiles from the Web of Science database, we analyze the gender differences in parenthood, academic achievements, and the mediation effect of work-family conflict and partner support in these gender gaps. Our results suggest that gender gaps in academic achievements are in fact “parenthood gaps”. Specifically, we found significant gender gaps exist in all measures of objective and subjective career achievements of academics in the parent group but not in the non-parent group. Additionally, mothers are more likely than fathers to experience higher levels of work-family conflict, and receive lower levels of partner support, contributing significantly to the gender gaps in objective and subjective career achievements for the parent group. Findings from this study identify the forms and the impact of parenthood on gender disparities in career achievements of academics and shed light on possible interventions and actions for mitigating gender inequalities in academia.

## Introduction

Gender disparity is prominent in academia. The science and engineering occupation, for example, is male-dominated (Ceci et al., 2014). In the U.S., women only have a share of 16% in science and engineering occupations in 2019 (National Center for Science and Engineering Statistics, 2021). Women are also disproportionally represented across all professorship ranks (Chesler et al., 2010; Mason et al., 2013), especially among full professors – only 32.5% of full professors in the U.S. are women in 2018 (Colby & Fowler, 2020). Studies show that, compared with men, women academics produce fewer papers (Larivière et al., 2013; Paul-Hus et al., 2015), receive fewer citations (Caplar et al., 2017; Maliniak et al., 2013), and have narrower collaboration networks (Ductor et al., 2021). Women are also less likely to receive project funding (Ley & Hamilton, 2008; Witteman et al., 2019) or prestigious awards (Lunnemann et al., 2019; Meho, 2021). These gendered differences are usually correlated and mutually reinforcing, contributing to women academics’ lower career satisfaction and higher attrition rates (Xu, 2008). Understanding variables and mechanisms underlying the gendered disadvantage for women is critical for achieving gender equality in academia.

Parenting is considered highly related to gender disparities in academia (Hunter & Leahey, 2010; Kelly & Grant, 2012; Morgan et al., 2021; Powell, 2021). It contributes to the gender gap in many key research performance metrics, such as scientific productivity (Mairesse et al., 2019; Morgan et al., 2021), citations (Lawson et al., 2021), and academic collaboration (Hunter & Leahey, 2010). There is also evidence that mother academics suffer from lower career satisfaction than father academics (Beckett et al., 2015), likely due to the reasons such as limited opportunities for tenure and promotion (Finkel & Olswang, 1996; Mason et al., 2013), lower salary (Kelly & Grant, 2012), and higher levels of work-family conflict (Martins et al., 2002). The prominent motherhood penalty could impel women to self-select away from academia and likely cause the academic pipeline leakage of women scientists from doctoral awardees to junior (and later senior) faculty (Anders, 2004). Yet, the present knowledge about parenting and gender disparities in academia has been shaped primarily by studies that are university-wide (e.g., Misra et al., 2012), monodisciplinary (e.g., Beckett et al., 2015), and thus of limited sample scale (e.g., Sonnert & Holton, 2020). The size, type, and mechanism of the association between parenting and academic achievements need further examination at a larger scale.

One prevailing reason parenting contributes to the gender disparity in academic careers is that women usually experience higher levels of work-family conflict than men due to parenting (Martins et al., 2002; van Daalen et al., 2006; Wang et al., 2020). Studies have indicated that women academics spend more time on childcare responsibilities and household tasks (Misra et al., 2012) and experience more research pressure and job stress than men (van Daalen et al., 2006). The multigenerational gender role beliefs, which characterize men and women with a simplistic set of features, may direct women academics toward women’s normative roles in the family (Ridgeway & Correll, 2004). Some women academics become “maternal gatekeepers” who are reluctant to relinquish family responsibility (Allen & Hawkins, 1999; Dickson, 2020; Hauser, 2012). Yet, limited research has examined how different forms of work-family contribute to the gender gap in academic career achievements.

The role of family support should also not be overlooked in gender differences in academic careers. Family has become “the newest battlefront in the struggle for gender equality” (Hauser, 2012), enduring and consequential for well-being across the life course (Thomas et al., 2017). Family support, especially partner support, represents a significant form of family-to-work enrichment and positive effects (Greenhaus & Powell, 2006; Kinnunen et al., 2006). Partner support can be instrumental or emotional (Adams & Golsch, 2021; Greenhaus & Powell, 2006) and may decrease work turnover rate and boost women’s perceptions of gains in productivity and career satisfaction (Ferguson et al., 2016; Juraqulova et al., 2015; Watanabe & Falci, 2016). Partner support can also relieve the tension inside the family and reduce the pressure from the family, mitigating work-family conflict and increasing well-being and career performance (Dickson, 2020; Ferguson et al., 2016; Thorstad et al., 2006; Wang et al., 2020). Although partner support is believed to be more critical to women’s career success than to men, especially for those in parenthood, women’s demand for partner support is less satisfied (Adams & Golsch, 2021; Dickson, 2020). Despite previous discussions on the role of partner support in family-to-work enrichment, limited attention has been paid to the relationship between partner support and the gender disparity in academia.

Using data from Web of Science (WoS) and 7,764 U.S. and Canada-based academics’ responses from our survey distributed in 2019 (see ***SI Appendix* Table S1-S5** for descriptive statistics), this study examines the gender gaps in academics’ objective and subjective career achievements and how parenting-related work-family conflict and partner support mediate these gaps. Objective career achievements are observable, socially recognized indicators signaling one’s human capital values (Valcour & Ladge, 2008). We use bibliometric indicators of scientific productivity, citation, and collaboration to measure objective career achievements (see ***Methods and Materials***). Subjective career achievements are one’s subjective feeling of career attainments (Ng et al., 2005), including self-reported research satisfaction, career satisfaction, and the perceived recognition by scholarly communities in this study.

Based on self-reported gender identification, the sample contains 4,425 (57.0%) women, 3,311 (42.7%) men, and 28 (0.4%) non-binaries. Due to the limited sample size for the non-binary group, our subsequent analyses only focus on women and men, which we admit is a limitation of our study. To better understand the role of parenting, we aggregated respondents based on their self-reported parenthood status: the parent group (*n*=5,670, 73.3%) and non-parent (*n*=1,534, 19.8%) group. Subsequent analyses refer to respondents in the parent group who self-identified as women and men as mothers and fathers, and those in the non-parent group as non-mothers and non-fathers. It should be noted that the two groups only include respondents who have ever married or cohabited (for two years or longer), given work-family conflict and partner support being variables of significant interest in this study.

## Results

### Parenthood and its overall compatibility with academic careers

Our results show significant gender differences in the parenthood status of academics. Across all disciplines, career stages, and races, a large majority (73.7%) of the respondents have at least one child. However, women (71.4%) are less likely than men (76.7%) to have children (OR=0.82, 95%CI [0.73, 0.91], p=0.000; see ***SI Appendix* Table S6**) and are more likely to have fewer children (Tobit coefficient=-0.14, 95%CI [−0.22, −0.07], p=0.000). This trend is different from the U.S. general population, where more adult women have children and have more children than men (Monte, 2017). Our results further show that the gender difference in parenthood status of academics is highly related to career considerations: For those with children, women are more likely than men to report that the number of children they have is related to career considerations (OR=2.34, 95%CI [2.08, 2.63], p= 0.000). Among those without children, women (59.9%) are more likely than men (43.2%) to report that they chose to be childfree due to career considerations (OR=2.10, 95%CI [1.69, 2.62], p=0.000; see ***SI Appendix* Table S7**), while controlling for discipline, career stage and race.

The gender difference in parenthood status in academia is likely due to the different perceptions of parenting compatibility with academic careers by women and men: Parenthood is perceived as less compatible with academic careers by women than by men (see **Table 1**). Women are more likely than men to report a negative impact on their career due to children (OR=0.46, 95%CI [0.41,0.51], p=0.000). More specifically, 71.3% of mothers reported negative child impact on their career, while only 48.6% of fathers indicated so. In contrast, 25.2% of fathers reported a positive impact on their career due to children, while only 14.1% of mothers indicated so. When aggregated by career stage, women are more likely to report more negative child impact during each career stage: trainee (OR=0.34, 95%CI [0.20,0.61], p=0.000), early career (OR=0.52, 95%CI [0.39,0.70], p=0.000), middle career (OR=0.47, 95%CI [0.40,0.56], p=0.000), and late career (OR=0.43, 95%CI [0.37,0.51], p=0.000).

**Table 1.**
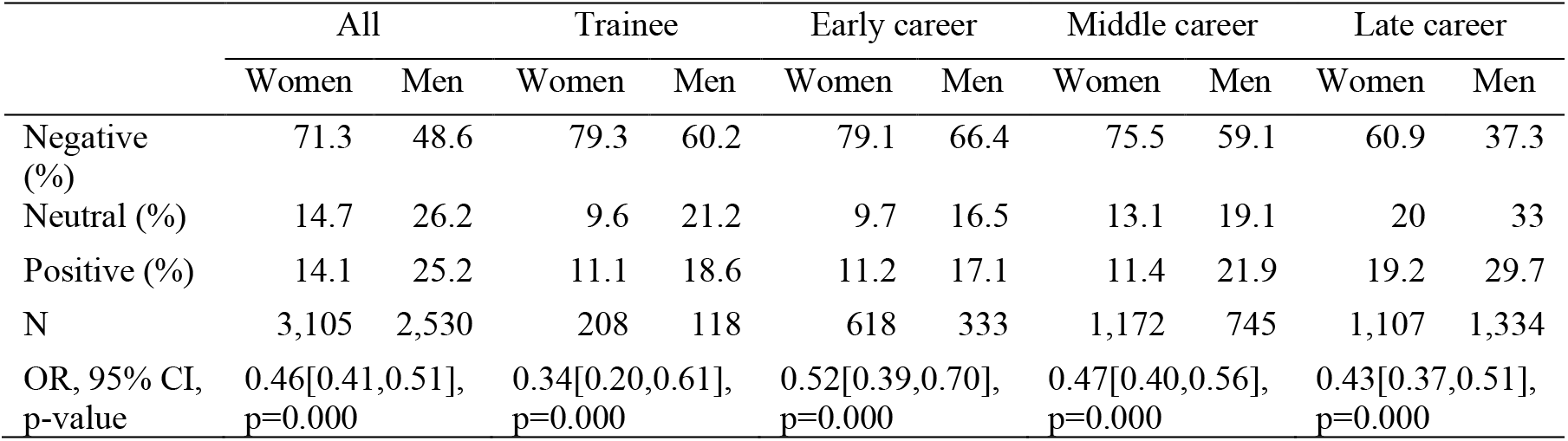
Child impact on career for parents. Odds ratio (women/men) values were based on ordinal logistic regression. Standard errors were clustered at the institution level. Control variables include discipline, race, and type of partner job.

### Gender gaps in subjective career achievements

We operationalized the measurement of subject career achievements using respondents’ self-reported satisfaction over research, career, and recognition by scholarly communities. Our results show that women academics, in general, are attached with lower levels of subjective career achievement than their male counterparts (see ***SI Appendix* Table S8**). Specifically, women are less likely than men to be satisfied with their research (OR= 0.76, 95%CI [0.68,0.85], p=0.000) and feel recognized by their scholarly communities (OR= 0.77, 95%CI [0.68,0.87], p=0.000).

Yet, the gendered differences vary by parenthood status. For the non-parent group, women and men show no significant differences in satisfaction over research, career, or the recognition by scholarly communities. For the parent group, however, the gender gap is prominent: Mother academics are less likely than fathers to be satisfied with their research achievements (OR= 0.72, 95%CI [0.63, 0.82], p=0.000), and the recognition from their scholarly communities (OR= 0.73, 95%CI [0.64, 0.84], p=0.000). While the satisfaction over research is significantly different between mothers and fathers, our results show no significant gender difference in the career satisfaction of parents. Research, teaching, and service being the three primary duties for many tenure-track faculty, it has been found that teaching and service are more heavily loaded on women than on men (Bellas, 1999; Guarino & Borden, 2017). Therefore, it is likely that women consider more on other gains (e.g., emotional gains) from teaching and service while assessing their overall career satisfaction, while men emphasize research. As shown in our results, of those women who expressed satisfaction over their career achievements, 13.6% were not satisfied with their research achievements. The number is 8.9% for men (see ***SI Appendix* Table S9**).

### Gender gaps in objective career achievements

We operationalized the measurement of objective career achievements for academics using publication-based indicators, given the “currency” role that scientific publications play in the academic reward system (Fogarty, 2009). The publication-based indicators used in this study include annual relative publication (ARP), average relative citation (ARC), and annual relative coauthor (ARCo). These indicators are used as measures for research productivity, citation, and the extent of collaboration, as normalized by discipline and time. The normalization was performed due to the varying scholarly practices across disciplines and the cumulative advantage in research achievements (such as citations) by time (see ***Materials and Methods***).

Our results further confirmed gender disparities in objective career achievements (See **Fig. 1** and ***SI Appendix* Table S10**). Regardless of parenthood status, women are outperformed by men in all three measures of objective career achievement. Specifically, women’s ARP is 0.28 units lower than men (95% CI [−0.39, −0.17], p=0.000), women’s ARC is 0.21 units lower than men (95% CI [−0.38,-0.05], p=0.013), and women’s ARCo is 0.08 unit lower than men (95% CI [−0.14,-0.01], p=0.023). This echoes previous findings that women produce fewer publications (Astegiano et al., 2019; Besselaar & Sandström, 2017), receive lower citations (Larivière et al., 2013; Maliniak et al., 2013), and have fewer coauthors (Ductor et al., 2021) than their male counterparts.

**Fig. 1.**
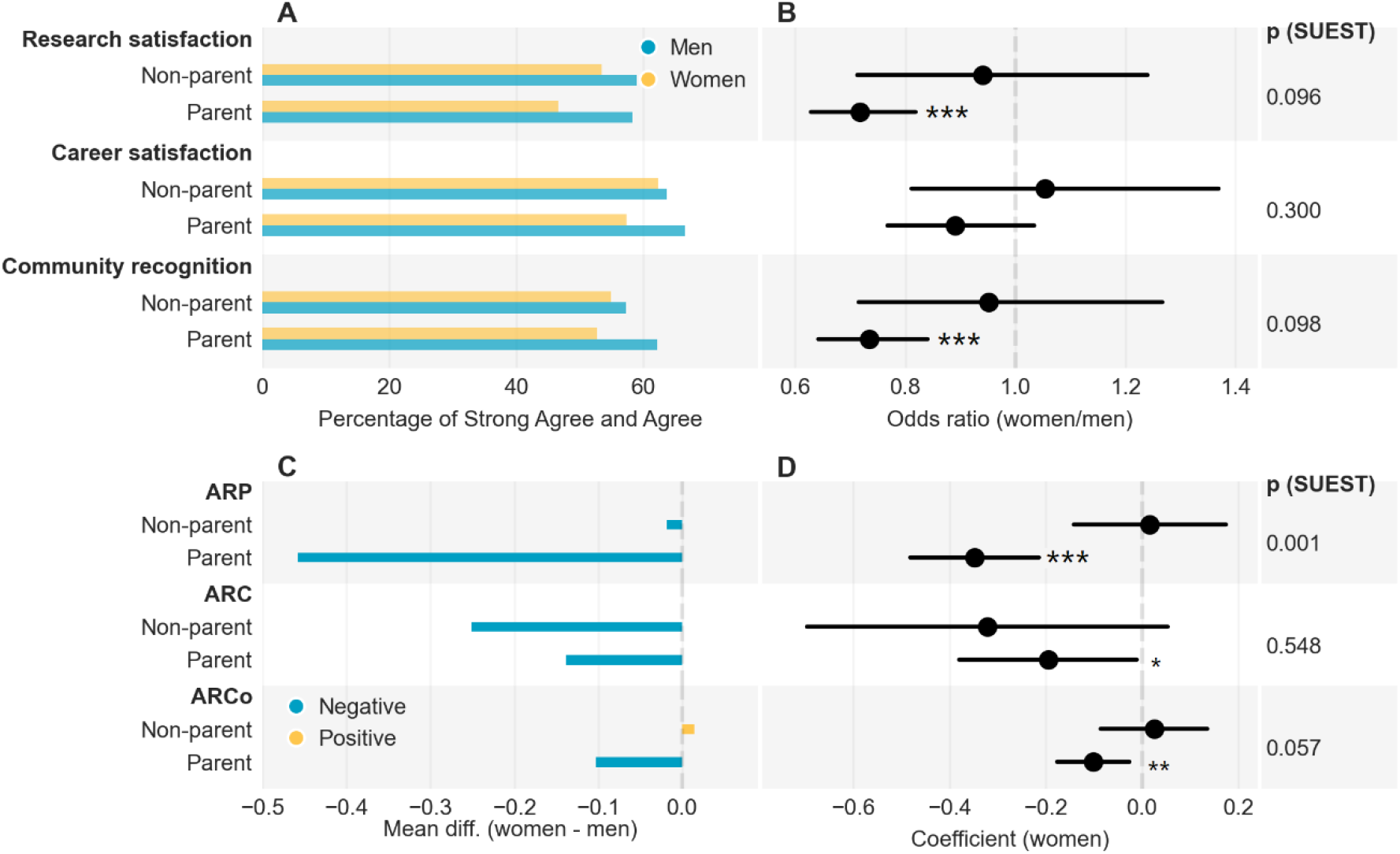
Subjective and objective career achievements by gender and parenthood status. **(A)** Percentage of satisfaction over research, career, and recognition by scholarly communities; (B) Women/men odds ratio for subjective career achievements; Control variables include discipline, career stage, partner job type, and race. Standard errors were clustered at the institution level. p(SUEST) values compare the odds ratio values between the parent and non-parent group; **(C)** Women-men difference in annual relative publication (ARP), average relative citation (ARC), and annual relative coauthor (ARCo). Positive values indicate female dominance and negative male dominance; (D) Coefficients for gender (women) based on linear regression analysis on ARP, ARC, and ARCo.

**Fig. 2.**
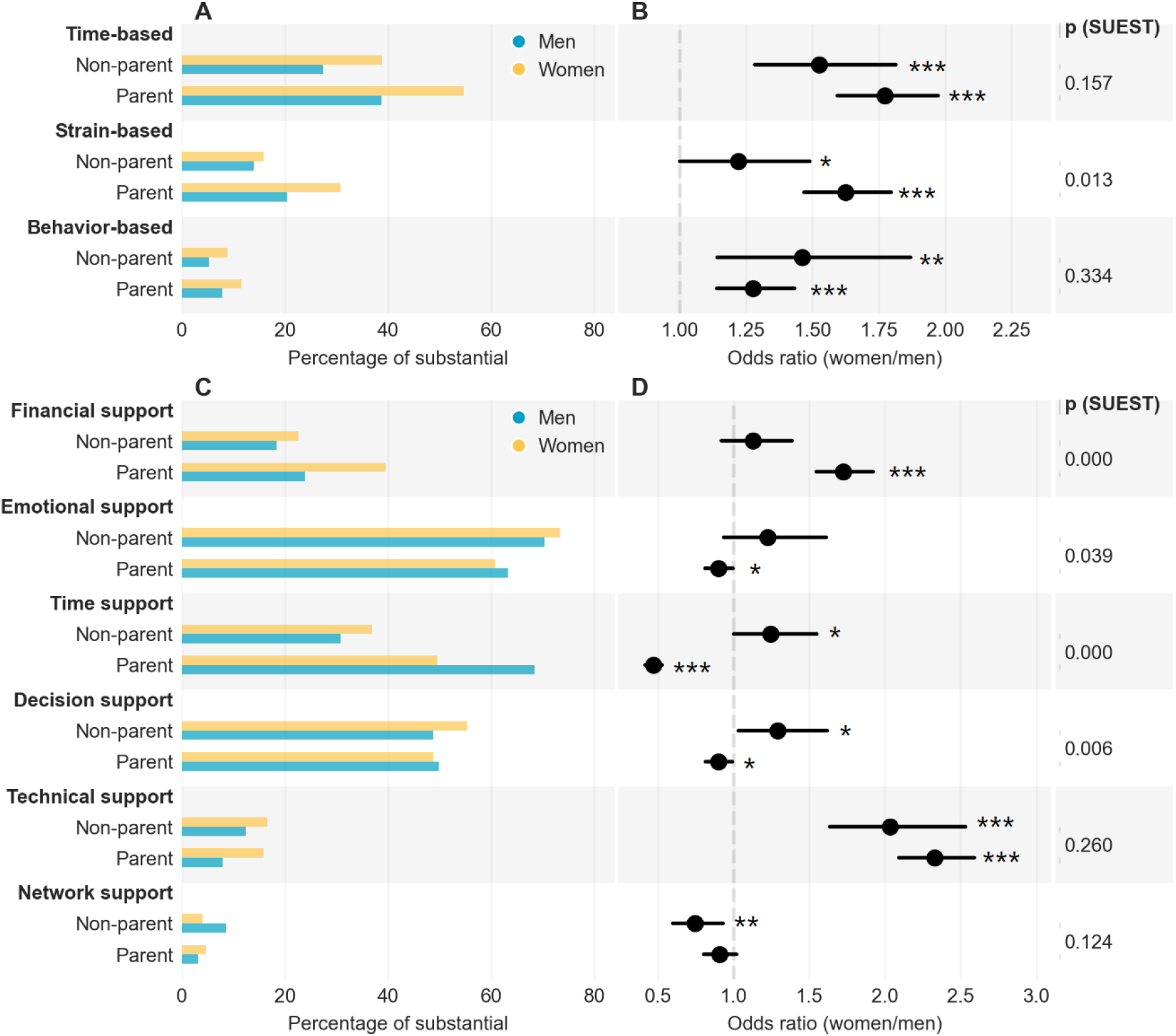
Forms of work-family conflict experienced, and partner support received by gender and parenthood status. (A) Percentage of women and men experiencing “substantial” conflict; (B) Odds ratio (women/men) for experiencing work-family conflict; (C) Percentage of women and men receiving “substantial” partner support; (D) Odds ratio (women/men) for receiving partner support. The (women/men) odds ratio values were based on logistic regression. Control variables include discipline, career stage, partner job type, and race. Standard errors were clustered at the institution level. SUEST was used to compare the odds ratio values between parent and non-parent group.

Yet, our results suggest parenthood may have played a significant role in the observed gender gaps in productivity, citation, and the extent of collaboration. In the parent group, mother academics’ ARP, ARC, and ARCo are 0.35 (95% CI [−0.48, −0.22], p=0.000), 0.20 (95% CI [−0.38, −0.01], p=0.038), and 0.10 units (95% CI [−0.18, −0.03], p=0.008) lower than that of father academics, respectively. However, in the non-parent group, women and men do not differ significantly in any of the three measures. Furthermore, the mean productivity difference between women and men is larger in the parent group than in the non-parent group: Our seemingly unrelated estimation (SUEST) analysis shows there is a significant difference between the gender coefficients (for ARP) of the parent and non-parent group regression analyses, indicating significant differences between productivity gender gaps between the parent and non-parent group (p=0.001). However, we do not observe such differences for citation and collaboration. These findings suggest that many previous findings on gender disparities in research productivity are likely the gender differences among parent academics, given that they are the majority of academics (73.7% of all academics in our study).

Moreover, recognizing that scholarly practices may vary as scholars advance through their careers, we further aggregated respondents by career stage. Results show that mother academics produced fewer publications than father academics across early, middle, and late careers. In the meantime, women and men in the non-parent group do not vary significantly in productivity during any career stage.

### Gender gaps in work-family conflict and partner support

When asked about factors that impeded their career, both genders agreed that work-family conflict is the major obstacle: 71.8% of women and 68.0% of men indicated they experienced “a little” to “substantial” levels of work-family conflict that impeded their career development, regardless of their parenthood status. However, the gender gap in work-family conflict is more significant in the parent group than in the non-parent group: Our results show mother academics are more likely than father academics to experience higher levels of work-family conflict (OR=1.31, 95%CI [1.19,1.45], p=0.000), but this does not hold for non-parent academics (OR=1.05, 95%CI= [0.85,1.28], p=0.662) (see ***SI Appendix* Table S11**).

Additionally, work-family conflict may occur in various forms related to childbearing and childrearing. Time-based conflict is the competition of time between different roles, e.g., mother vs. researcher; strain-based conflict happens when the strain in one role interferes with one’s ability to perform in another role; behavior-based conflict happens when the behavior requirements as a role become incompatible with the behavioral expectation of another (Greenhaus & Beutell, 1985). Our results show that mothers are more likely than fathers to experience higher levels of time-based (OR=1.77, 95%CI [1.59,1.97], p=0.000), strain-based (OR=1.62, 95%CI [1.47,1.80], p=0.000), and behavior-based (OR=1.28, 95%CI [1.14,1.43], p=0.000) work-family conflict.

On the other hand, we should not overlook the role of partner support, which is beneficial for career success in many areas. We asked about the levels of partner support for career received by academics, including financial, emotional, time, decision, technical, and network support. Our results show that, women academics are overall more likely than men to receive higher levels of financial (OR=1.55, 95%CI [1.41, 1.71], p=0.000) and technical support (OR= 2.28, 95%CI [2.06, 2.53], p=0.000), but less like to receive higher levels of time (OR= 0.58, 95%CI [0.52, 0.64], p=0.000) and network support (OR= 0.87, 95%CI [0.78, 0.97], p=0.012) from their partners. Yet, the gender difference in partner support, similar to that in work-family conflict, also varies by parenthood status. Mothers are less likely than fathers to receive higher levels of time support (OR= 0.47, 95%CI [0.42, 0.53], p=0.000), a pattern that does not hold in the non-parent group. Additionally, mothers are more likely than fathers to receive more financial support (OR=1.72, 95%CI [1.55, 1.92], p=0.000), while non-mothers are not (OR=1.13, 95%CI [0.92,1.39], p=0.248). Mothers are also less likely than fathers to receive higher levels of decision support (OR= 0.90, 95%CI [0.82, 1.00], p=0.040), and less likely to receive higher levels of emotional support (OR=0.90, 95%CI [0.81,1.00], p=0.046). Women are more likely than men to receive higher levels of technical support in both the parent (OR=2.33, 95%CI [2.09,2.59], p=0.000) and non-parent (OR=2.03, 95%CI [1.63,2.53], p=0.000) group (see **Fig. 3** and ***SI Appendix* Table S12**), which aligns with the finding that women are usually disproportionally associated with technical work in research (Macaluso et al., 2016). We further found that the gender difference in received support varies significantly by respondents’ parenthood status: The women to men odds ratios for the parent group vary significantly from that for the non-parent group regarding financial (p=0.000), emotional (p=0.039), time (p=0.000), and decision support (p=0.006).

**Fig. 3.**
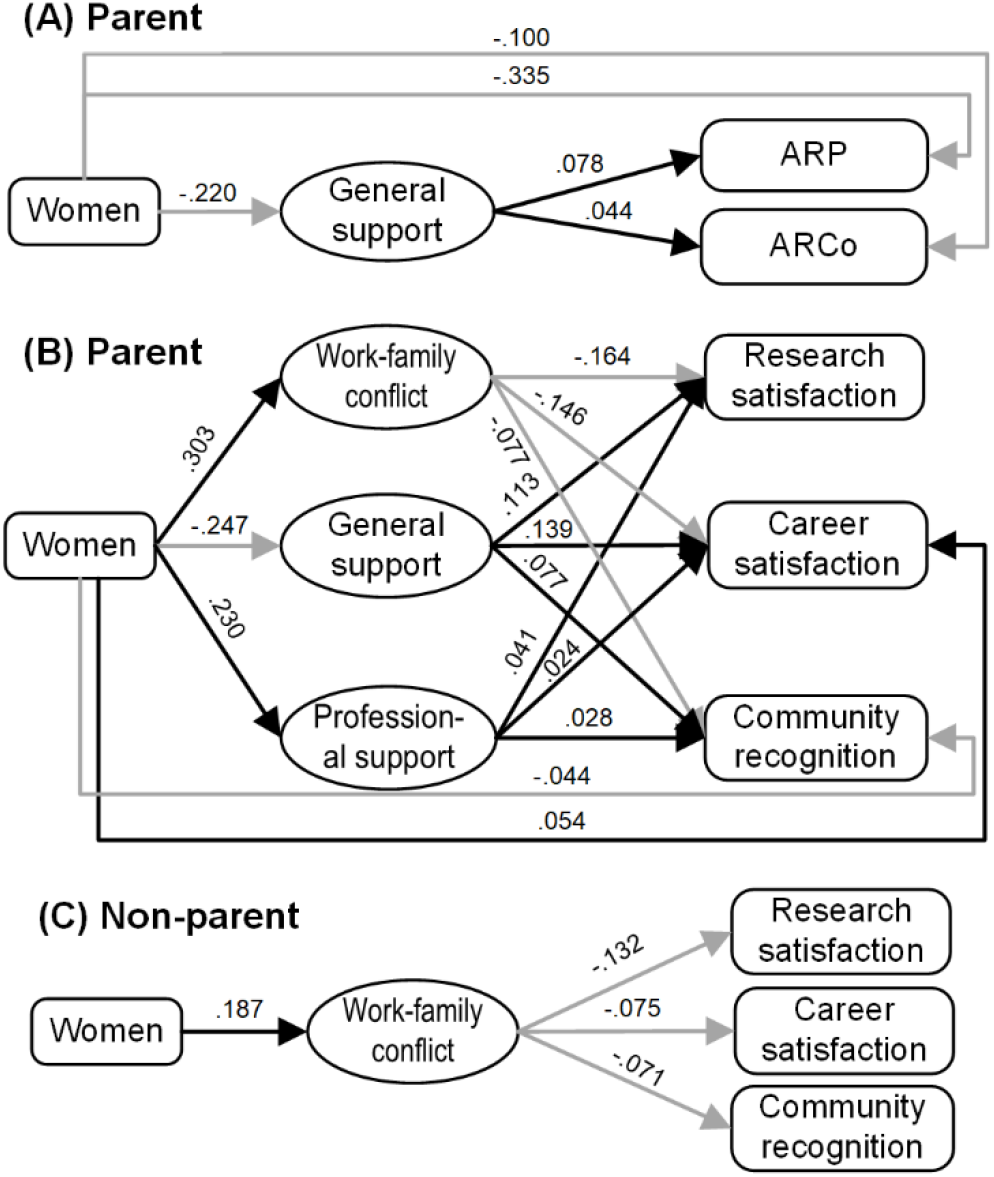
Mediation effect analysis models of partner support and work-family conflict between gender and subjective and objective career achievement measures. Only paths with statistically significant effects (p < 0.05) are shown. Black and gray lines denote positive and negative mediating coefficients, respectively. **(A)** sets the objective career achievement as the outcomes and tests with the parent group (n=4,173). **(B)** sets the subjective career achievement measures as the outcomes and tests with the parent group (n=4,557). **(C)** sets the subjective career achievement measures as the outcomes and tests with the non-parent group (n=1,152).

### The mediation effect of work-family conflict and partner support

While findings above suggest the gender gaps in academics’ objective and subjective career achievements, it is unclear what roles work-family conflict and partner support play in forming these gaps. We then used mediation effect analysis to unveil the possible mediating role of work-family conflict and partner support. To increase the interpretability of variables, we first extracted four factors from the three types of work-family conflict and six forms of partner support (nine variables in total) based on principal component analysis and varimax rotation, with each attached with at least one variable contributing significantly (factor loading ≥ 0.5) to the factor. The four factors explained 66.69% of the total variance. The labels for the four factors are: work-family conflict (including time-, strain- and behavior-based conflict), financial support, professional support (including technical and network support), and general support (including emotional, decision, and time support, all of which do not require special skills or economic power) (see ***SI Appendix* Table S13**).

The mediation effect analysis based on the extracted factors shows that work-family conflict and partner support are significant mediator variables contributing to the association between gender and objective and subjective career achievement measures for parents, albeit with different underlying mechanisms (see **Fig. 4** and ***SI Appendix* Table S14**). General support is a mediating variable for the association between gender and parents’ objective career achievement. Mothers received less general support than fathers, which contributes significantly to mothers’ lower levels of annual relative productivity (Path effect [PE]=-0.017, 95% percentile CI [−0.031, −0.004]) and fewer annual relative unique coauthors (PE=-0.010, 95% percentile CI [−0.018, −0.002]). For the association between gender and subjective career achievement measures, mothers are subject to higher levels of work-family conflict than fathers, which leads to lower levels of research satisfaction (PE=-0.050, 95% percentile CI [−0.062, −0.038]) and scholarly community recognition (PE =-0.023, 95% percentile CI [−0.031, −0.016]). The significantly less general support received by mothers are also shown to disadvantage mothers in research satisfaction, career satisfaction, and scholarly community recognition (PE=-0.028, 95% percentile CI [−0.037, −0.019]; PE=-0.034, 95% percentile CI [−0.045, −0.025]; PE=-0.019, 95% percentile CI [−0.026, −0.013]). On the contrary, mothers receive more professional support, which mitigates the negative association between gender and the three measures, albeit with comparatively small indirect effects (PE=0.009, 95% percentile CI [0.003, 0.016]; PE=0.006, 95% percentile CI [0.000, 0.011]; PE=0.006, 95% percentile CI [0.001, 0.012]). For non-parent academics, the mediating effect of work-family conflict is also observed between the association of gender and subjective career success: women are subject to higher levels of work-family conflict than men, inhibiting women’s satisfaction over research satisfaction (PE=-0.025, 95% percentile CI [−0.045, −0.008]), career satisfaction (PE=-0.014, 95% percentile CI [−0.028, −0.004]), and scholarly community recognition (PE=-0.013, 95% percentile CI [−0.026, −0.003]). Work-family conflict and partner support show no significant mediation effect on the associations between gender and the objective career achievement for the non-parent group.

## Discussion

Our analysis of the parenthood status of academics provides new insights into the dilemma many women academics face: motherhood or academic career. On the one hand, mother academics are significantly more likely than fathers to report higher levels of negative impact on career due to children, reflecting the ‘motherhood penalty’ on academic career (Bonache et al., 2022; Misra et al., 2012). The perceived negative impact of parenting is one of the major systematic barriers for women’s self-selecting attrition from academia, contributing to the “leaky pipeline” in academia (Anders, 2004). On the other hand, we found that women are less likely than men to have children and have fewer children. Women in academia who reach the same career achievements as their male counterparts tend to slough off the burden of children, which may restrict their working time, networking opportunities, and career development opportunities. Given the motherhood penalty, it is unsurprising that many women stay in academia by paying the price of being childfree or having fewer children than men: Our results show non-mothers are more likely than non-fathers to report they chose to be childfree due to career considerations and mother academics are more likely than father academics to report that the lower number of children they have is due to career considerations.

Our study also unveils gender gaps in both objective and subjective career achievements of academics and suggests much of the gender gap only exists in the parent group. Specifically, we find gender differences in all measures of subjective career achievement and objective career achievement for the parent group. For the non-parent group, however, no significant gender difference was found in any career achievement measures. We thus argue that given many previous studies (Bendels et al., 2018; Caplar et al., 2017; Holliday et al., 2014; Larivière et al., 2013) reported overall gender disparities in academia without differentiating between parents and non-parents, these disparities may primarily derive from the differences between academic mothers and fathers. Follow-up studies on gender disparities could further examine this issue by comparing parents and non-parents.

Given the role of productivity in the academic reward system, current policies and regulations that aim to provide short-term support for childcaring and recovery right after birth or adoption are not sufficient to bridge the gender gap in academia. Furthermore, the impact of parenting on the research output of mothers may last longer than asserted by previous studies (Morgan et al., 2021): We find significant gender differences in productivity across all career stages among the parents, but no such differences have been observed among the non-parents. This suggests that policies and interventions addressing gender inequalities in the scientific workforce should go beyond short-term assistance such as parental leaves and extend long-term support for mothers. Other forms of sustainable family-friendly support, such as subsidized childcare, onsite childcare, flexible working schedules, and supportive working environments, are available options (Feeney & Stritch, 2019).

Our findings on mothers’ higher chances of encountering more elevated levels of work-family conflict in all the forms of time-, strain-, and behavior-based conflict confirm that mothers indeed suffer more from multiple social roles: mother and researcher. Previous findings suggested that mothers are the primary caregiver of children, which typically requires extra time and engagement in parenting activities (Dickson, 2020). We also found that mothers are significantly more likely than fathers to experience time-based conflict and less likely to receive the time, emotional, and decision support from partners, a dangerous signal for the family to sustain marital satisfaction and gender equality (Mickelson et al., 2006; Thorstad et al., 2006). On the contrary, mothers are more likely to receive financial and technical support from their partners. It reflects the entrenched gender division of household labor that expands with the new childcare task: Mothers are recognized, and even morally self-recognized, as the babysitters and fathers are the breadwinners (Hauser, 2012). As a social norm, it is more common that mothers sacrifice their working time, emotionally support fathers’ work, and understand their decisions because fathers bear the financial responsibility. In contrast, fathers compensate mothers’ loss due to extra childcare responsibility by offering financial and technical support.

Our mediating effect analysis results provide new evidence for the claim that family is “the newest battlefront in the struggle for gender equality” (Hauser, 2012) for academia, confirming that family-related reasons are significantly associated with gender gaps in career achievements of parents. Specifically, mother academics receive less general support (including time, emotional and decisional support) from their partners than father academics, contributing to the gender gap in productivity and collaboration. It corroborates the finding that shortened working time and the limited decision support for more work engagement restrict mothers’ opportunities for networking and academic collaboration (Finkel & Olswang, 1996; Mason et al., 2013). The limited general support from partners thus may hurt mother academics’ research productivity directly or indirectly through constraining research collaborations, a factor that usually leads to higher productivity (Abbasi et al., 2011; Sonnenwald, 2007). Moreover, the higher levels of work-family conflict experienced by mothers also worsen the gender gap in satisfaction over research and perceived recognition by parents, which are highly related to women’s turnover intention (Watanabe & Falci, 2016). Reducing mothers’ work-family conflict and increasing their general support could thus reduce the gender gap in academics’ perceived career achievement and the high attrition rates of mothers in the scientific workforce due to lower satisfaction rates. These mediating variables indicate the directions for mitigating the negative impacts of motherhood: Partner support in the forms of time, emotional, and decision support, and other collective efforts to reduce the time-based, strain-based, and behavior-based conflict for mothers.

By combining large-scale survey and bibliometric data, we reveal the decisive role of parenthood in the current gender gap in academia and underlying mechanisms on how parenting-related conflicts and support mediate the gap. Previous studies asserted the importance of various forms of support from governments and institutions (Morgan et al., 2021). Yet, our results show that family is another new battleground: assistance and support provided by partners in time, emotional and decisional support are also vital. Addressing gender inequality in academia is a task requiring collective intelligence. In addition to parenting-related support by governments and institutions, which could be sometime inadequate due to restrictions of resources and regulations in the U.S., the support from partners and families will also help narrow the gender gap in academia.

## Limitations

Like many other survey-based studies, our research is not immune to potential bias caused by the self-selection of respondents. Given the topic of our survey, it is not surprising that a higher proportion of women than men in the population may have responded (see ***SI Appendix* Table S1**). Future efforts on this topic should consider strategies in encouraging more responses from men academics to ensure the representativeness of both genders. We also restricted our survey to be sent to individuals associated with institutions in the U.S. or Canada only, considering the similarity between the “laddered” academic systems between the two countries and their distinctions from other countries and regions. But this omitted the rising research power in other countries and areas and restricted the generalizability of our findings to the scientific workforce in North America. Future research may examine this issue in other countries and regions of the world to understand the gender gap in the global scientific workforce.

Furthermore, we operationalized objective career achievement in productivity, citation, and the extent of collaboration as metrics of publication, citation, and coauthor. Yet, publication practices of academics only account for a part of their career achievements, and other measures such as grant funding and prestigious award have been overlooked. Moreover, it is known that disciplines relying on venues other than journals are underrepresented in WoS, given that WoS largely indexes journal publications. Therefore, we may underestimate respondents’ productivity, citation, and collaboration within those disciplines. The strategy we used to normalize these measures by field helps mitigate this issue, given we calculated respondents’ measures of productivity, citation, and collaboration proportional to fellow researchers in the same disciplines. Finally, the cross-sectional questionnaire data from our study restricted the potential to explore time-series change. To analyze the causal relationship of parenting on academics’ career achievements, we plan to track the respondents and organize similar surveys in the future, comparing the parents as the treatment group with non-parents as the control group within gender.

## Materials and Methods

### Data Collection

This study relies on two data sources: a large-scale survey distributed to 99,168 researchers and their publication profiles from the WoS database by Clarivate Analytics. For the survey, we extracted from WoS 396,674 researchers who published at least one paper from 2000 to 2019, were affiliated with an institution located in the U.S. or Canada, and had a valid email address associated with them. We then randomly sampled 99,168 researchers (25%) from the population and sent a survey with 53 questions about parenting and career development in 2019 through Qualtrics (see ***SI Appendix Survey questions***). A total of 10,333 respondents initiated the survey, of which 9,105 finished. An analysis of the attrition failed to identify a common point of departure, suggesting individual variability in dropout rather than failed survey construction. This study’s final number of respondents is **7,764**, after removing respondents lacking information for critical variables of interest (see ***SI Appendix Survey procedures***). We also collected data for non-binary academics, who were excluded from our analysis due to insufficient sample (***n***=28) (See ***SI Appendix* Table S2-S5**).

To assess the objective career achievement of respondents, we extracted the bibliographic records for individuals, including productivity (for which our proxy was papers), citation (for which our proxy was citation), and extent of collaboration (for which our proxy as unique coauthors). Recognizing the difference in the publication practices across disciplines and the cumulative nature of these measures, we normalized the three indicators of objective career achievement by discipline and time (see ***SI Appendix Objective career achievement measures***). We then use the annual relative productivity (ARP), average relative citation (ARC), and annual relative coauthor (ARCo) as indicators for productivity, citation, and extent of collaboration.

### Statistical Analysis

This study used binary logistic regression, ordinal logistic regression, linear regression, and Tobit regression model to explore the gender difference in academic careers, depending on the measurement scale of outcome variables. In logistic regression analysis, the odds ratio (OR) of gender (women over men) is computed to explore the role of gender in outcome variables. An OR value lower than 1 indicates women are less likely to produce the outcome than men. Linear regression analysis yields a coefficient for gender (women=1 and men=0) rather than odds ratio, where a value below (above) 0 indicates that being a woman has a negative (positive) impact on the outcome measure. We used Tobit regression model to estimate the relationship between parents’ gender and child number, which is censored as the survey took six as a threshold for child number.

A consistent list of variables was used as control variables across the study, including disciplinary area, career stage, partner job type, and race. We did not include the institution as one control variable because the respondents’ current affiliated institution may also be the outcome of their previous publications and career satisfaction in other workplaces. Instead, we clustered the standard errors in regressions at the institution level, considering an unobserved part of residual may be correlated within an institution (e.g., an institution created a high-intensity work culture that influenced its employees’ publishing behaviors satisfaction). We also used seemingly unrelated estimation (SUEST) to compare if gender differences vary significantly across parents and non-parents (see ***SI Appendix* Statistical analysis**). Observations with missing values were removed without imputation.

This study analyzed the mediating effect of work-family conflict and partner support in the association between gender and career achievements of academics using mediation effect analysis. Before performing the analysis, we first extracted four factors from partner support and work-family conflict variables using principal component analysis and varimax rotation. The Cronbach’s alpha of each factor’s principal variables is > 0.6, showing their acceptable internal consistency within a factor (see ***SI Appendix* Table S13**). We constructed a mediation effect analysis model for all subjective and objective career achievement measures to test the indirect effects mediated by the four factors from gender to the outcome measures for the parent group (see ***SI Appendix* Table S14**). A bootstrap sampling procedure was used with 5,000 iterations to compute 95% confidence intervals and statistical significances for all indirect effect paths—the estimation controlled for career stage, disciplinary area, race, and partner job type. Standard errors are clustered by academics’ affiliations to account for the non-independence of observations in the same affiliation.

## Data availability

All data needed to evaluate the conclusions in the paper are present here and the Supplementary Materials. Aggregated or de-identified data on variables used in this study is available on GitHub (https://github.com/UWMadisonMetaScience/parenting).

## Acknowledgments

Support for this research was provided by the Office of the Vice Chancellor for Research and Graduate Education at the University of Wisconsin–Madison with funding from the Wisconsin Alumni Research Foundation. We thank Observatoire des sciences et des technologies at the University of Quebec in Montreal for access to the Web of Science data.

