## Supplemental Information for "The gender gap in academic career achievements and the mediation effect of work-family conflict and partner support"

**This PDF file includes:**

Supplementary Information Text
Tables S1 to S14
Survey Questions
SI References

### Supplementary Information Text

#### Survey procedures

We constructed the population and sample of the survey from an in-house version of the Web of Science (WoS) database by Clarivate Analytics. To do so, we first extracted 396,674 researchers (population) who published at least one paper between 2000 and 2019, had a valid email address attached to them, and were affiliated with an institution in US or Canada. We restricted our population to US and Canada based on the consideration that these two countries have a similarly “laddered” system for academic career advancement, which is different from some other countries and regions. We then randomly sampled 99,168 (25% of the population) academics and sent a questionnaire with 53 questions about family and academic career development in September 2019 through email using Qualtrics survey software (see ***Survey Questions***). Among those surveyed, 10,333 were initiated, and 9,105 completed the survey. An analysis of the attrition failed to identify a common point of departure, suggesting individual variability in dropout rather than failed survey construction. Additional 202 responses were removed due to the missing answers to critical questions of interest, e.g., partnership status, parenthood status, and gender (see **Table S1** and **Table S2**). We also excluded respondents if their self-identified rank or role is in the *student, lecturer, technician or technical assistant,* and *others*. These roles are generally not considered research-oriented, an essential factor in this study. Furthermore, given partner support being a variable of interest in this study, we only included those who are or were married or cohabited for two years or longer in the final analytical sample, containing 7,764 respondents.

#### Operationalization of key variables

**Gender.** Gender is of the primary explanatory variables of interest in this study. The self-identified gender category was used to assign the gender for individuals, which includes women, men, and non-binary. Because we were only able to collect 28 responses in the non-binary gender category, we excluded the category from regression and mediation effect analysis but provided the descriptive statistics for it (see **Table S2**)

**Partnership and parenthood status.** This study defines a marriage or a domestic relationship of 2 years or longer as partnership. This study further categorized the partnership status of respondents as ever married or cohabited (including now and before) and never married or cohabited based on their responses to a relative question in the survey. As stated above, the never-married-or-cohabited group was excluded in this analysis. We classified the parenthood status of respondents based on their responses to the question asking for the number of children (includes step-, adopted, and biological children of all ages) they have. Those who reported having 0 child constitutes the non-parent group, and the rest the parent group (see **Table S3** for statistics).

**Control variables.** While performing regression analyses, including the regressions in the mediation effect analysis, we controlled for some variables that may affect the outcome variables. The list of control variables includes disciplinary area, career stage, race, and partner job type. The disciplinary area includes arts & humanities, medical sciences, natural science & engineering, and social sciences. Respondents were assigned to a disciplinary area based on self-reported data from the survey (see **Table S2**). The career stage of individuals was decided based on the self-identified rank/role information from the survey. There are four levels of career stage used in our analysis: trainee (including post-doctoral fellow and research associate in the survey), early career (assistant professor in the survey), middle career (including associate professor and senior researcher in the survey), and late career (including full and emeritus professor in the survey) (see **Table S3**). Another variable we controlled for in our analyses is race. Based on survey responses, there are two race categories: white (including white) and non-white (including Black or African American; American Indian or Native American; Asian or Pacific Islander; Hispanic or Latino; and others). We also controlled for the partner job types (research-oriented or not research-oriented) identified by respondents in the survey.

**Objective career achievement measures**

We developed three indicators based on publication to assess the objective career achievement of academics: Annual relative publication, average relative, and annual relative coauthors. These measures are discipline- and time-based normalizations, given that publication practices usually vary by discipline and are of cumulative advantage. The disciplines are assigned by the classification developed for the National Science Foundation, which classifies each journal into one discipline and one specialty. The details of each indicator are as follows:

**a) Annual relative publication (ARP).** We used ARP, the number of publications normalized by discipline and years that a respondent has been publishing based on WoS, to represent productivity. It is intended to measure the annual productivity of a researcher in relation to fellow researchers in the same discipline. The discipline of each scholar was decided based on the discipline of publications they authored as indexed by WoS. ARP is calculated as follows:

For one academic $x$, we first compute their yearly productivity (YP) by dividing the total number of publications using the year span between x's newest and oldest publication:

$$YP_x = \frac{total\ publications}{latest\ year - earliest\ year + 1}$$

where $total\ publications$ is the total number of papers authored by academic $x$, $latest\ year$ is the year of their latest publication, and $earliest\ year$ is the year of their earliest publication.

The academic $x$'s relative publication is

$$RP_x = YP_x \Big/ \frac{1}{n} \sum_{i=1}^{n} YP_i$$

where $YP_i$ is the yearly publications research $i$ in a discipline by WoS, and $n$ is the total number of academics in the discipline.

**b) Average relative citation (ARC).** ARC is the discipline- and time-based citation count in relation to fellow researchers in the same discipline for papers published in the same year. For each discipline and each year in WoS, we compute an index of base citation (BC) as

$$BC = \frac{1}{m} \sum_{k=1}^{m} citation_k$$

where $citation_k$ is the citations received by the $k$th paper in that discipline published in that year, and $m$ is the total number of papers in that discipline published in that year. The ARC of academic $x$ is defined as:

$$ARC_x = \frac{1}{n} \sum_{i=1}^{n} \frac{citation_i}{BC_i}$$

where $citation_i$ is the citations received by the $i$th paper by academic $x$, $BC_i$ is the base citations of the discipline and year in which the $i$th paper was published, and $n$ is the total number of papers by academic $x$.

**c) Annual relative coauthor (ARCo).** ARCo measures researchers' extent of collaboration using the number of unique coauthors they collaborated with annually in relation to fellow researchers in the same discipline. We first compute academic $x$'s yearly unique coauthors (YUC):

$$YUC_x = \frac{total\ unique\ coauthors}{latest\ year - earliest\ year + 1}$$

where $total\ unique\ coauthor$ is the number of unique coauthors in the byline of papers by academic $x$, $latest\ year$ is the year of their latest publication, and $earliest\ year$ is the year of their earliest publication. The academic $x$'s ARCO is

$$ARCO_x = YUC_x / \frac{1}{n} \sum_{i=1}^{n} YUC_i$$

where $YUC_i$ is the yearly unique authors of the $i$th academic in the discipline, and $n$ is the total number of academics in that discipline. It should be noted that papers with more than 100 authors in their bylines were excluded from the calculation (about 2.1% of papers by our respondents) to avoid possible distortions caused by "hyper-authorship" in some disciplines (1).

#### Statistical analysis

**Regression analysis.** We used several regression analysis techniques to explore the gendered difference in academic careers, including logistic regression, ordinal logistic regression, Tobit model (censored normal regression), and linear regression. Specific procedures and analysis methods vary by the scale of dependent variables. Ordinal logistic regressions were used for dependent variables of ordinal scales (e.g., Likert scale questions), regular logistic regressions were used for dichotomous dependent variables (e.g., yes/no questions), and linear regression were used for continuous dependent variables (e.g., counts). Tobit model is used for the dependent variable of child number, which is censored at the upper threshold of 6. The robust standard errors of all the above regression models are clustered by respondents' affiliated institutions, which is identified by the respondents' email domains, to address the intra-institution correlation.

**Seemingly unrelated estimation.** We used seemingly unrelated estimation (SUEST) to compare if gendered differences in academic careers vary significantly by the parenthood status of academics. SUEST is a technique for comparing the coefficients of different regression analyses whose covariances are non-zero, including the ones used in the present study (2, 3). For example, in the analysis of gender and childcare impact on research development, we use SUEST to test if the odds ratio of women over men differs between the non-parent and parent groups. For group $i$, we first build a benchmark model that for every option $j$,

$$\log\frac{P(y_i \le j)}{P(y_i > j)} = \beta_{i0} + \beta_{i1}Gender_i + \beta_{i2}Role_i + \beta_{i3}Area_i + \beta_{i4}Race_i + \beta_{i5}PartnerCareer_i$$

To test if $\beta_{11} = \beta_{21}$, we "stack" the two group samples into a stacked dataset with the number of observations equal to the two groups' sum and estimate the following equation

$$\log\frac{P(y_{stacked} \le j)}{P(y_{stacked} > j)} = \gamma_{10} + \gamma_{11}Gender1 + \gamma_{12}Role1 + \gamma_{13}Area1 + \gamma_{14}Race1 + \gamma_{15}PartnerCareer1 + \gamma_{20} + \gamma_{21}Gender2 + \gamma_{22}Role2 + \gamma_{23}Area2 + \gamma_{24}Race2 + \gamma_{25}PartnerCareer2$$

Where each observation keeps each original independent variable value in a new independent variable named by a combination of the original variable name and the observation's group number. The other independent variable values are fixed to 0. For example, $Gender1$ keeps the data in $Gender_1$ for the observations in the non-parent group and fills in 0 for the observations in the parent group. The vector of $y_{stacked}$ is $\begin{bmatrix} y_1 \\ y_2 \end{bmatrix}$. The standard errors are clustered by each respondent's institution. After estimation, a Wald test is performed to test if $\gamma_{11}$ equals to $\gamma_{21}$ and compute its statistical significance, for which details are shown by Clogg et al. and Mize et al. (2, 3).

### Supplementary Tables

**Table S1. Gender composition of the population, surveyed, respondents, and analytical sample.** The gender categorization was estimated using the methods successfully applied in Larivière et al. (4). This is only used for a rough assessment of the percentage of each gender in the population, sample, responses, and the final analytical sample. Self-identified gender information was used to assign the gender in all other analyses of this study.

| Gender | Population | | Surveyed | | Respondents | | Analytical Sample | |
|---|---|---|---|---|---|---|---|---|
| | N | % | N | % | N | % | N | % |
| Men | 189,046 | 47.3 | 44,725 | 45.10 | 4,126 | 39.9 | 3,075 | 39.6 |
| Women | 113,507 | 28.4 | 29,850 | 30.10 | 4,003 | 38.7 | 3,043 | 39.2 |
| Unknown | 97,121 | 24.3 | 24,594 | 24.80 | 2,204 | 21.3 | 1,646 | 21.2 |
| All | 396,674 | 100 | 99,168 | 100 | 10,333 | 100 | 7,764 | 100 |

**Table S2. Sample distribution by gender, career stage and disciplinary area.**

| | Trainee | | Early Career | | Middle Career | | Late Career | | Total | |
|---|---|---|---|---|---|---|---|---|---|---|
| | N | % | N | % | N | % | N | % | N | % |
| **Natural Science & Engineering** | | | | | | | | | | |
| Women | 180 | 48.5 | 220 | 54.9 | 371 | 46.7 | 407 | 39.2 | 1,178 | 45.2 |
| Men | 188 | 50.7 | 178 | 44.4 | 420 | 52.9 | 631 | 60.7 | 1,417 | 54.4 |
| Non-binary | 3 | 0.8 | 3 | 0.7 | 3 | 0.4 | 1 | 0.1 | 10 | 0.4 |
| All | 371 | 100.0 | 401 | 100.0 | 794 | 100.0 | 1,039 | 100.0 | 2,605 | 100.0 |
| **Medical Sciences** | | | | | | | | | | |
| Women | 195 | 70.4 | 417 | 69.4 | 555 | 66.8 | 472 | 45.4 | 1,639 | 62.9 |
| Men | 80 | 28.9 | 178 | 29.6 | 273 | 32.9 | 426 | 41.0 | 957 | 36.7 |
| Non-binary | 2 | 0.7 | 6 | 1.0 | 3 | 0.4 | 0 | 0.0 | 11 | 0.4 |
| All | 277 | 100.0 | 601 | 100.0 | 831 | 100.0 | 898 | | 2,607 | 100.0 |
| **Social Sciences** | | | | | | | | | | |
| Women | 99 | 76.2 | 293 | 69.3 | 495 | 70.4 | 355 | 34.2 | 1,242 | 63.6 |
| Men | 30 | 23.1 | 128 | 30.3 | 206 | 29.3 | 341 | 32.8 | 705 | 36.1 |
| Non-binary | 1 | 0.8 | 2 | 0.5 | 2 | 0.3 | 0 | 0.0 | 5 | 0.3 |
| All | 130 | 100.0 | 423 | 100.0 | 703 | 100.0 | 696 | | 1,952 | 100.0 |
| **Arts & Humanities** | | | | | | | | | | |
| Women | 7 | 50.0 | 59 | 71.1 | 139 | 65.6 | 161 | 55.3 | 366 | 61.0 |
| Men | 7 | 50.0 | 23 | 27.7 | 72 | 34.0 | 130 | 44.7 | 232 | 38.7 |
| Non-binary | 0 | 0.0 | 1 | 1.2 | 1 | 0.5 | 0 | 0.0 | 2 | 0.3 |
| All | 14 | 100.0 | 83 | 100.0 | 212 | 100.0 | 291 | 100.0 | 600 | 100.0 |
| **All areas** | | | | | | | | | | |
| Women | 481 | 60.7 | 989 | 65.6 | 1,560 | 61.4 | 1,395 | 47.7 | 4,425 | 57.0 |
| Men | 305 | 38.5 | 507 | 33.6 | 971 | 38.2 | 1,528 | 52.3 | 3,311 | 42.6 |
| Non-binary | 6 | 0.8 | 12 | 0.8 | 9 | 0.4 | 1 | 0.0 | 28 | 0.4 |
| All | 792 | 100.0 | 1,508 | 100.0 | 2,540 | 100.0 | 2,924 | 100.0 | 7,764 | 100.0 |

#### Table S3. Sample distribution by career stage, role, and gender

| Stage | Role | Women | Men | Non-binary | Total |
|---|---|---|---|---|---|
| Trainee | Post-doctoral fellow | 291 | 184 | 4 | 479 |
| | Research associate | 190 | 121 | 2 | 313 |
| Early Career | Assistant professor | 989 | 507 | 12 | 1,508 |
| Middle Career | Associate professor | 1,350 | 762 | 7 | 2,119 |
| | Senior researcher | 210 | 209 | 2 | 421 |
| Late Career | Full professor | 1,216 | 1,216 | 1 | 2,433 |
| | Emeritus professor | 179 | 312 | 0 | 491 |
| All | | 4,425 | 3,311 | 28 | 7,764 |

#### Table S4. Sample distribution by parenthood status, gender and career stage

| Career Stage | Women | | | | Men | | | | Non-binary | |
|---|---|---|---|---|---|---|---|---|---|---|
| | Parent | | Non-parent | | Parent | | Non-parent | | Parent | Non-parent |
| | N | % | N | % | N | % | N | % | N | N |
| Trainee | 208 | 53.2 | 183 | 46.8 | 118 | 49.2 | 122 | 50.8 | 0 | 2 |
| Early Career | 626 | 70.3 | 265 | 29.7 | 336 | 71.8 | 132 | 28.2 | 2 | 6 |
| Middle Career | 1,183 | 81.5 | 268 | 18.5 | 747 | 81.1 | 174 | 18.9 | 4 | 3 |
| Late Career | 1,115 | 82.8 | 231 | 17.2 | 1,337 | 89.4 | 159 | 10.6 | 1 | 0 |
| Total | 3,132 | 76.8 | 947 | 23.2 | 2,538 | 81.2 | 587 | 18.8 | 7 | 11 |

#### Table S5. Summary of objective career achievement measures by WoS discipline

| WoS discipline | Number of respondents | Mean of yearly productivity | Mean of yearly unique coauthors |
|---|---|---|---|
| Arts | 48 | 1.01 | 1.44 |
| Biology | 559 | 1.07 | 4.67 |
| Biomedical Research | 581 | 1.09 | 6.50 |
| Chemistry | 214 | 1.12 | 4.98 |
| Clinical Medicine | 1495 | 1.14 | 6.41 |
| Earth and Space | 477 | 1.10 | 5.51 |
| Engineering and Technology | 379 | 1.08 | 4.28 |
| Health | 339 | 1.07 | 4.72 |
| Humanities | 303 | 1.00 | 1.15 |
| Mathematics | 207 | 1.04 | 2.44 |
| Physics | 191 | 1.26 | 5.54 |
| Professional Fields | 714 | 1.01 | 2.26 |
| Psychology | 577 | 1.05 | 3.46 |
| Social Sciences | 955 | 1.00 | 1.93 |
| Other | 725 | / | / |

**Table S6. Odds ratio of parenthood status by gender.** The odds ratio (OR) of gender was computed using logistic regression to measure women's relative odds of having children over men's odds of doing so. Control variables include disciplinary area, career stage, and race. Standard errors have been clustered at the institution level.

| | **Women** | | **Men** | | **Both** | | **OR, 95% CI, p-value** |
|---|---|---|---|---|---|---|---|
| | **N** | **%** | **N** | **%** | **N** | **%** | |
| Parent | 3,160 | 71.4 | 2,540 | 76.7 | 5,700 | 73.7 | 0.82[0.73,0.91], p= 0.000 |
| Non-parent | 1,265 | 28.6 | 771 | 23.3 | 2,036 | 26.3 | |

**Table S7. The Number of children related to career considerations.** The odds ratios (OR) of gender are computed by ordinal logistic regression to measure women's relative odds of answering a more positive option over men's odds. Control variables include area, career stage, partner job type, and race. Standard errors have been clustered at the institution level. The original answers have been recategorized into Negative, Neutral, and Positive.

| | **Non-parent** | | | **Parent** | | | **Both** | | |
|---|---|---|---|---|---|---|---|---|---|
| | **Women** | **Men** | **Both** | **Women** | **Men** | **Both** | **Women** | **Men** | **Both** |
| No | 37.1 | 56.8 | 46.5 | 49.3 | 70.9 | 59.0 | 47.2 | 68.3 | 56.4 |
| Yes | 60.0 | 43.2 | 53.5 | 50.7 | 29.1 | 41.0 | 52.8 | 31.7 | 43.6 |
| N | 926 | 579 | 1,505 | 3,091 | 2,477 | 5,599 | 4,017 | 3,087 | 7,104 |
| OR, 95% CI, p-value | 2.10 [1.66,2.67], p=0.000 | | | 2.34 [2.08, 2.63], p=0.000 | | | 2.29 [2.08, 2.53], p=0.000 | | |

**Table S8. Satisfaction over research and career, and recognition by scholarly communities.** The odds ratios (OR) of gender are computed by ordinal logistic regression to measure women's relative odds of answering a higher level of agreement over men's odds. Control variables include area, career stage, partner job type, and race. Standard errors have been clustered at the institution level. The original answers have been recategorized as Disagree, Neutral, and Agree.

| | Non-parent | | | Parent | | | Both | | |
|---|---|---|---|---|---|---|---|---|---|
| | **Women** | **Men** | **Both** | **Women** | **Men** | **Both** | **Women** | **Men** | **Both** |
| **Research satisfaction** | | | | | | | | | |
| Disagree (%) | 21.2 | 19.2 | 20.4 | 28.6 | 20.8 | 25.1 | 26.9 | 20.5 | 24.1 |
| Neutral (%) | 2.6 | 3.2 | 2.8 | 3.1 | 2.5 | 2.8 | 3.0 | 2.6 | 2.8 |
| Agree (%) | 76.3 | 77.6 | 76.8 | 68.3 | 76.8 | 72.1 | 70.1 | 76.9 | 73.1 |
| N | 931 | 563 | 1,494 | 3,098 | 2,496 | 5,594 | 4,029 | 3,059 | 7,088 |
| OR, 95% CI, p-value | 0.94[0.71,1.24], p=0.662 | | | 0.72[0.63,0.82], p=0.000 | | | 0.76[0.68,0.85], p=0.000 | | |
| **Career satisfaction** | | | | | | | | | |
| Disagree (%) | 15.2 | 14.7 | 15.0 | 18.5 | 14.6 | 16.8 | 17.7 | 14.6 | 16.4 |
| Neutral (%) | 2.2 | 3.9 | 2.8 | 3.4 | 2.9 | 3.1 | 3.1 | 3.1 | 3.1 |
| Agree (%) | 82.7 | 81.4 | 82.2 | 78.1 | 82.5 | 80.1 | 79.2 | 82.3 | 80.5 |
| N | 928 | 559 | 1,487 | 3,097 | 2,500 | 5,597 | 4,025 | 3,059 | 7,084 |
| OR, 95% CI, p-value | 1.05[0.81,1.37], p=0.694 | | | 0.89[0.77,1.03], p=0.129 | | | 0.91[0.81,1.04], p=0.158 | | |
| **Scholarly recognition** | | | | | | | | | |
| Disagree (%) | 11.2 | 11.3 | 11.3 | 15.0 | 9.8 | 12.7 | 14.2 | 10.1 | 12.4 |
| Neutral (%) | 6.8 | 6.1 | 6.5 | 7.0 | 5.8 | 6.4 | 6.9 | 5.8 | 6.4 |
| Agree (%) | 82.0 | 82.7 | 82.2 | 78.0 | 84.4 | 80.9 | 78.9 | 84.1 | 81.2 |
| N | 925 | 560 | 1,485 | 3,090 | 2,489 | 5,579 | 4,015 | 3,049 | 7,064 |
| OR, 95% CI, p-value | 0.95[0.71,1.27], p=0.735 | | | 0.73[0.64,0.84], p=0.000 | | | 0.77[0.68,0.87], p=0.000 | | |

**Table S9. Summary of respondents who agree with they are satisfied with career progress but disagree with they are satisfied with research progress.** The % column denotes the percentage of the respondents who fit the same conditions and agree that they are satisfied with career progress.

| | Men | | Women | | Both | |
|---|---|---|---|---|---|---|
| | **N** | **%** | **N** | **%** | **N** | **%** |
| Non-parent | 38 | 8.0 | 83 | 10.7 | 121 | 9.7 |
| Parent | 189 | 9.1 | 355 | 14.6 | 544 | 12.0 |
| Both | 227 | 8.9 | 438 | 13.6 | 665 | 11.5 |

**Table S10. Annual relative publication, average relative citation, and annual relative coauthors by gender, parenthood status, and career stage.** Regression coefficients of gender were computed by multiple linear regression to measure the average differences between women and men. Control variables include area, career stage, partner job type, and race. Standard errors have been clustered at the institution level.

| Career stage | Gender | Non-parent | | Parent | | Both | |
|---|---|---|---|---|---|---|---|
| | | **Mean** | **n** | **Mean** | **n** | **Mean** | **n** |
| **Annual Relative publication** | | | | | | | |
| All | Women | 1.70 | 838 | 1.82 | 2822 | 1.79 | 3660 |
| | Men | 1.71 | 515 | 2.27 | 2285 | 2.17 | 2800 |
| | Both | 1.70 | 1353 | 2.02 | 5107 | 1.96 | 6460 |
| | Coefficient, 95% CI, p-value | 0.02[-0.14,0.17], p=0.843 | | -0.35[-0.48,-0.22], p=0.000 | | -0.28[-0.39,-0.17], p=0.000 | |
| Trainee | Women | 1.65 | 161 | 1.47 | 192 | 1.55 | 353 |
| | Men | 1.45 | 104 | 1.87 | 103 | 1.66 | 207 |
| | Both | 1.57 | 265 | 1.61 | 295 | 1.59 | 560 |
| | Coefficient, 95% CI, p-value | 0.13[-0.15,0.42], p=0.352 | | -0.43[-0.87,0.00], p=0.052 | | -0.05[-0.20,0.10], p=0.527 | |
| Early Career | Women | 1.66 | 233 | 1.77 | 561 | 1.74 | 794 |
| | Men | 1.71 | 118 | 2.00 | 302 | 1.92 | 420 |
| | Both | 1.67 | 351 | 1.85 | 863 | 1.80 | 1214 |
| | Coefficient, 95% CI, p-value | -0.03[-0.28,0.22], p=0.805 | | -0.24[-0.44,-0.03], p=0.023 | | -0.08[-0.20,0.04], p=0.203 | |
| Middle Career | Women | 1.57 | 237 | 1.63 | 1050 | 1.62 | 1287 |
| | Men | 1.66 | 149 | 1.94 | 664 | 1.89 | 813 |
| | Both | 1.60 | 386 | 1.75 | 1714 | 1.72 | 2100 |
| | Coefficient, 95% CI, p-value | -0.00[-0.29,0.29], p=0.989 | | -0.26[-0.42,-0.10], p=0.001 | | -0.13[-0.25,-0.01], p=0.028 | |
| Late Career | Women | 1.93 | 207 | 2.10 | 1019 | 2.07 | 1226 |
| | Men | 1.95 | 144 | 2.56 | 1216 | 2.49 | 1360 |
| | Both | 1.94 | 351 | 2.35 | 2235 | 2.29 | 2586 |
| | Coefficient, 95% CI, p-value | -0.06[-0.50,0.37], p=0.776 | | -0.46[-0.68,-0.23], p=0.000 | | -0.28[-0.43,-0.14], p=0.000 | |
| **Average relative citation** | | | | | | | |
| All | Women | 1.93 | 838 | 2.13 | 2822 | 2.08 | 3660 |
| | Men | 2.22 | 515 | 2.28 | 2285 | 2.27 | 2800 |
| | Both | 2.04 | 1353 | 2.20 | 5107 | 2.16 | 6460 |
| | Coefficient, 95% CI, p-value | -0.32[-0.70,0.05], p=0.093 | | -0.20[-0.38,-0.01], p=0.038 | | -0.21[-0.38,-0.05], p=0.013 | |
| Trainee | Women | 1.42 | 161 | 1.86 | 192 | 1.66 | 353 |
| | Men | 1.91 | 104 | 1.93 | 103 | 1.92 | 207 |
| | Both | 1.61 | 265 | 1.88 | 295 | 1.75 | 560 |
| | Coefficient, 95% CI, p-value | -0.55[-1.02,-0.07], p=0.026 | | -0.13[-0.62,0.35], p=0.590 | | -0.26[-0.58,0.06], p=0.110 | |
| Early Career | Women | 1.99 | 233 | 1.98 | 561 | 1.98 | 794 |
| | Men | 2.39 | 118 | 1.97 | 302 | 2.09 | 420 |
| | Both | 2.12 | 351 | 1.98 | 863 | 2.02 | 1214 |
| | Coefficient, 95% CI, p-value | -0.42[-1.35,0.51], p=0.373 | | -0.07[-0.38,0.24], p=0.649 | | -0.21[-0.48,0.05], p=0.110 | |
| Middle Career | Women | 2.00 | 237 | 2.04 | 1050 | 2.03 | 1287 |
| | Men | 1.80 | 149 | 1.95 | 664 | 1.93 | 813 |
| | Both | 1.92 | 386 | 2.00 | 1714 | 1.99 | 2100 |
| | Coefficient, 95% CI, p-value | 0.13[-0.41,0.67], p=0.628 | | 0.06[-0.25,0.36], p=0.719 | | -0.11[-0.37,0.15], p=0.411 | |

| Career stage | Gender | Non-parent | | Parent | | Both | |
|---|---|---|---|---|---|---|---|
| | | **Mean** | **n** | **Mean** | **n** | **Mean** | **n** |
| Late Career | Women | 2.17 | 207 | 2.36 | 1019 | 2.33 | 1226 |
| | Men | 2.72 | 144 | 2.57 | 1216 | 2.58 | 1360 |
| | Both | 2.40 | 351 | 2.47 | 2235 | 2.46 | 2586 |
| | Coefficient, 95% CI, p-value | -0.52[-1.35,0.30], p=0.212 | | -0.43[-0.75,-0.10], p=0.010 | | -0.37[-0.59,-0.14], p=0.002 | |
| **Annual relative coauthor** | | | | | | | |
| All | Women | 0.97 | 838 | 0.98 | 2822 | 0.98 | 3660 |
| | Men | 0.95 | 515 | 1.09 | 2285 | 1.06 | 2800 |
| | Both | 0.96 | 1353 | 1.03 | 5107 | 1.01 | 6460 |
| | Coefficient, 95% CI, p-value | 0.02[-0.09,0.14], p=0.661 | | -0.10[-0.18,-0.03], p=0.008 | | -0.08[-0.14,-0.01], p=0.023 | |
| Trainee | Women | 1.13 | 161 | 1.07 | 192 | 1.10 | 353 |
| | Men | 0.96 | 104 | 1.37 | 103 | 1.16 | 207 |
| | Both | 1.06 | 265 | 1.18 | 295 | 1.12 | 560 |
| | Coefficient, 95% CI, p-value | 0.18[-0.09,0.46], p=0.188 | | -0.32[-0.78,0.13], p=0.165 | | -0.03[-0.15,0.10], p=0.684 | |
| Early Career | Women | 1.02 | 233 | 1.10 | 561 | 1.08 | 794 |
| | Men | 1.08 | 118 | 1.19 | 302 | 1.16 | 420 |
| | Both | 1.04 | 351 | 1.13 | 863 | 1.11 | 1214 |
| | Coefficient, 95% CI, p-value | -0.04[-0.24,0.16], p=0.703 | | -0.08[-0.23,0.07], p=0.307 | | -0.01[-0.11,0.08], p=0.757 | |
| Middle Career | Women | 0.91 | 237 | 0.91 | 1050 | 0.91 | 1287 |
| | Men | 0.93 | 149 | 1.04 | 664 | 1.02 | 813 |
| | Both | 0.91 | 386 | 0.96 | 1714 | 0.95 | 2100 |
| | Coefficient, 95% CI, p-value | 0.03[-0.24,0.31], p=0.828 | | -0.12[-0.23,0.00], p=0.053 | | -0.05[-0.13,0.03], p=0.243 | |
| Late Career | Women | 0.84 | 207 | 0.98 | 1019 | 0.96 | 1226 |
| | Men | 0.87 | 144 | 1.06 | 1216 | 1.04 | 1360 |
| | Both | 0.85 | 351 | 1.02 | 2235 | 1.00 | 2586 |
| | Coefficient, 95% CI, p-value | -0.04[-0.28,0.20], p=0.755 | | -0.07[-0.18,0.03], p=0.170 | | -0.04[-0.12,0.04], p=0.318 | |

**Table S11. Work-family conflict and its forms by gender and parenthood status.** The odds ratios (OR) of gender were computed by ordinal logistic regression to measure women's relative odds of answering a higher degree of option over men's odds. Control variables include area, career stage, partner job type, and race. Standard errors have been clustered at the institution level.

| | **Non-parents** | | | **Parents** | | | **Both** | | |
|---|---|---|---|---|---|---|---|---|---|
| | **Women** | **Men** | **Both** | **Women** | **Men** | **Both** | **Women** | **Men** | **Both** |
| **Work-family conflict** | | | | | | | | | |
| Not at all (%) | 52.0 | 53.0 | 52.4 | 21.3 | 27.4 | 24.0 | 28.2 | 32.0 | 29.8 |
| A little (%) | 29.1 | 28.3 | 28.8 | 30.5 | 33.4 | 31.8 | 30.2 | 32.5 | 31.2 |
| Moderate (%) | 12.0 | 11.9 | 12.0 | 25.6 | 24.4 | 25.1 | 22.6 | 22.1 | 22.4 |
| Substantial (%) | 6.9 | 6.9 | 6.9 | 22.6 | 14.9 | 19.2 | 19.1 | 13.4 | 16.7 |
| N | 883 | 540 | 1,423 | 3,061 | 2,460 | 5,521 | 3,944 | 3,000 | 6,944 |
| OR, 95% CI, p-value | 1.05[0.85,1.28], p=0.662 | | | 1.31[1.19,1.45], p=0.000 | | | 1.21[1.11,1.31], p=0.000 | | |
| **Time-based conflict** | | | | | | | | | |
| Not at all (%) | 9.5 | 14.0 | 11.2 | 2.1 | 5.9 | 3.8 | 3.8 | 7.4 | 5.4 |
| A little (%) | 22.1 | 24.8 | 23.1 | 14.0 | 20.1 | 16.7 | 15.8 | 21.0 | 18.1 |
| Moderate (%) | 29.4 | 34.1 | 31.2 | 29.2 | 34.9 | 31.8 | 29.2 | 34.8 | 31.6 |
| Substantial (%) | 39.1 | 27.1 | 34.6 | 54.7 | 39.0 | 47.7 | 51.2 | 36.8 | 45.0 |
| N | 919 | 557 | 1,476 | 3,091 | 2,484 | 5,575 | 4,010 | 3,041 | 7,051 |
| OR, 95% CI, p-value | 1.53[1.28,1.82], p=0.000 | | | 1.77[1.59,1.97], p=0.000 | | | 1.68[1.53,1.84], p=0.000 | | |
| **Strain-based conflict** | | | | | | | | | |
| Not at all (%) | 28.5 | 31.4 | 29.6 | 10.6 | 18.2 | 14.0 | 14.7 | 20.6 | 17.2 |
| A little (%) | 31.8 | 34.8 | 32.9 | 28.0 | 32.8 | 30.2 | 28.9 | 33.2 | 30.7 |
| Moderate (%) | 23.7 | 19.8 | 22.2 | 30.7 | 28.4 | 29.7 | 29.1 | 26.9 | 28.1 |
| Substantial (%) | 16.1 | 14.1 | 15.3 | 30.8 | 20.6 | 26.2 | 27.4 | 19.4 | 23.9 |
| N | 916 | 555 | 1,471 | 3,083 | 2,479 | 5,562 | 3,999 | 3,034 | 7,033 |
| OR, 95% CI, p-value | 1.22[1.00,1.49], p=0.049 | | | 1.62[1.47,1.80], p=0.000 | | | 1.48[1.36,1.62], p=0.000 | | |
| **Behavior-based conflict** | | | | | | | | | |
| Not at all (%) | 63.8 | 71.4 | 66.7 | 57.1 | 61.7 | 59.2 | 58.7 | 63.5 | 60.7 |
| A little (%) | 16.4 | 13.8 | 15.4 | 17.9 | 18.8 | 18.3 | 17.5 | 17.8 | 17.7 |
| Moderate (%) | 11.0 | 9.8 | 10.5 | 13.5 | 11.7 | 12.7 | 12.9 | 11.3 | 12.2 |
| Substantial (%) | 8.8 | 5.1 | 7.4 | 11.5 | 7.9 | 9.9 | 10.9 | 7.4 | 9.4 |
| N | 909 | 552 | 1,461 | 3,063 | 2,459 | 5,522 | 3,972 | 3,011 | 6,983 |
| OR, 95% CI, p-value | 1.46[1.14,1.87], p=0.003 | | | 1.28[1.14,1.43], p=0.000 | | | 1.29[1.17,1.43], p=0.000 | | |

**Table S12. Level of partner support by gender and parenthood status.** The odds ratios (OR) of gender were computed by ordinal logistic regression to measure women's relative odds of answering a higher degree of option over men's odds. Control variables include area, career stage, partner job type, and race. Standard errors have been clustered at the institution level.

| | Non-parents | | | Parents | | | Both | | |
|---|---|---|---|---|---|---|---|---|---|
| | **Women** | **Men** | **Both** | **Women** | **Men** | **Both** | **Women** | **Men** | **Both** |
| **Financial support** | | | | | | | | | |
| Not at all (%) | 30.5 | 30.1 | 30.4 | 16.5 | 24.1 | 19.9 | 19.8 | 25.2 | 22.1 |
| A little (%) | 23.8 | 30.5 | 26.3 | 20.1 | 27.1 | 23.2 | 20.9 | 27.7 | 23.9 |
| Moderate (%) | 22.8 | 21.1 | 22.2 | 23.8 | 24.7 | 24.2 | 23.6 | 24.0 | 23.8 |
| Substantial (%) | 22.8 | 18.4 | 21.1 | 39.6 | 24.1 | 32.6 | 35.7 | 23.0 | 30.2 |
| N | 898 | 545 | 1,443 | 3,000 | 2,437 | 5,437 | 3,898 | 2,982 | 6,880 |
| OR, 95% CI, p-value | 1.13[0.92,1.39], p=0.248 | | | 1.72[1.55,1.92], p=0.000 | | | 1.55[1.41,1.71], p=0.000 | | |
| **Emotional support** | | | | | | | | | |
| Not at all (%) | 1.2 | 2.0 | 1.5 | 3.2 | 2.4 | 2.8 | 2.7 | 2.3 | 2.5 |
| A little (%) | 7.9 | 7.9 | 7.9 | 12.4 | 11.7 | 12.1 | 11.4 | 11.0 | 11.2 |
| Moderate (%) | 17.0 | 19.9 | 18.1 | 23.6 | 22.7 | 23.2 | 22.1 | 22.2 | 22.1 |
| Substantial (%) | 73.9 | 70.2 | 72.5 | 60.8 | 63.3 | 61.9 | 63.8 | 64.5 | 64.1 |
| N | 923 | 554 | 1,477 | 3,078 | 2,469 | 5,547 | 4,001 | 3,023 | 7,024 |
| OR, 95% CI, p-value | 1.23[0.93,1.61], p=0.141 | | | 0.90[0.81,1.00], p=0.046 | | | 0.96[0.87,1.05], p=0.373 | | |
| **Time support** | | | | | | | | | |
| Not at all (%) | 6.4 | 8.0 | 7.0 | 4.0 | 2.6 | 3.4 | 4.5 | 3.6 | 4.1 |
| A little (%) | 20.9 | 22.8 | 21.6 | 18.5 | 8.3 | 14.0 | 19.0 | 10.9 | 15.6 |
| Moderate (%) | 35.4 | 38.6 | 36.6 | 27.9 | 20.6 | 24.7 | 29.6 | 23.8 | 27.1 |
| Substantial (%) | 37.3 | 30.6 | 34.8 | 49.6 | 68.4 | 58.0 | 46.9 | 61.7 | 53.2 |
| N | 874 | 526 | 1,400 | 3,044 | 2,411 | 5,455 | 3,918 | 2,937 | 6,855 |
| OR, 95% CI, p-value | 1.25[1.00,1.55], p=0.049 | | | 0.47[0.42,0.53], p=0.000 | | | 0.58[0.52,0.64], p=0.000 | | |
| **Decision support** | | | | | | | | | |
| Not at all (%) | 8.6 | 9.0 | 8.7 | 11.8 | 8.5 | 10.3 | 11.1 | 8.6 | 10.0 |
| A little (%) | 12.4 | 16.4 | 13.9 | 15.0 | 14.0 | 14.5 | 14.4 | 14.4 | 14.4 |
| Moderate (%) | 23.3 | 26.0 | 24.3 | 24.4 | 27.9 | 26.0 | 24.2 | 27.6 | 25.7 |
| Substantial (%) | 55.8 | 48.6 | 53.0 | 48.8 | 49.7 | 49.2 | 50.4 | 49.5 | 50.0 |
| N | 841 | 523 | 1,364 | 2,775 | 2,315 | 5,090 | 3,616 | 2,838 | 6,454 |
| OR, 95% CI, p-value | 1.29[1.03,1.62], p=0.026 | | | 0.90[0.82,1.00], p=0.040 | | | 0.97[0.89,1.06], p=0.545 | | |
| **Technical support** | | | | | | | | | |
| Not at all (%) | 24.2 | 39.1 | 29.8 | 26.8 | 46.4 | 35.6 | 26.2 | 45.1 | 34.3 |
| A little (%) | 34.3 | 30.9 | 33.1 | 34.1 | 30.9 | 32.7 | 34.1 | 30.9 | 32.7 |
| Moderate (%) | 24.9 | 17.9 | 22.3 | 23.3 | 14.6 | 19.4 | 23.7 | 15.2 | 20.0 |
| Substantial (%) | 16.7 | 12.0 | 14.9 | 15.8 | 8.1 | 12.4 | 16.0 | 8.8 | 12.9 |
| N | 877 | 524 | 1,401 | 2,893 | 2,332 | 5,225 | 3,770 | 2,856 | 6,626 |
| OR, 95% CI, p-value | 2.03[1.63,2.53], p=0.000 | | | 2.33[2.09,2.59], p=0.000 | | | 2.28[2.06,2.53], p=0.000 | | |
| **Network support** | | | | | | | | | |
| Not at all (%) | 53.9 | 46.5 | 51.1 | 56.8 | 54.4 | 55.7 | 56.2 | 53.0 | 54.8 |
| A little (%) | 27.3 | 29.7 | 28.2 | 27.6 | 30.0 | 28.7 | 27.5 | 30.0 | 28.6 |
| Moderate (%) | 14.6 | 15.1 | 14.8 | 10.8 | 12.3 | 11.5 | 11.7 | 12.8 | 12.2 |
| Substantial (%) | 4.2 | 8.8 | 5.9 | 4.8 | 3.2 | 4.1 | 4.6 | 4.2 | 4.5 |
| N | 840 | 525 | 1,365 | 2,823 | 2,333 | 5,156 | 3,663 | 2,858 | 6,521 |
| OR, 95% CI, p-value | 0.75[0.60,0.93], p=0.010 | | | 0.91[0.81,1.02], p=0.110 | | | 0.87[0.78,0.97], p=0.012 | | |

#### Table S13. Factor analysis and internal consistency measure result of partner support and work-family conflict.

| Items | Factor loading | | | | Commu-nality | Cronbach's alpha |
|---|---|---|---|---|---|---|
| | Work-family Conflict | General support | Professional support | Financial support | | |
| Time conflict | 0.78 | | | | 0.63 | 0.65 |
| Strain conflict | 0.82 | | | | 0.68 | |
| Behavior conflict | 0.69 | | | | 0.51 | |
| Emotional support | | 0.71 | | | 0.58 | 0.62 |
| Time support | | 0.78 | | | 0.62 | |
| Decision support | | 0.74 | | | 0.58 | |
| Technical support | | | 0.83 | | 0.71 | 0.63 |
| Network support | | | 0.84 | | 0.72 | |
| Financial support | | | | 0.98 | 0.97 | |
| Variance Explained | 19.63% | 18.92% | 16.82% | 11.32% | | |

**Table S14.** Mediation effect analysis results for subjective and objective career achievement measures

| Mediator | Outcome | a (gender → mediator) | p-value | b (mediator → outcome) | p-value | ab (path effect) or c' (direct effect, gender → outcome), Bootstrap standard error | 95% Percentile CI | |
|---|---|---|---|---|---|---|---|---|
| | | | | | | | Lower bound | Upper bound |
| Subjective career success | | | | | | | | |
| Non-parents | | | | | | | | |
| (Direct effect) | Research satisfaction | | | | | 0.000(0.052) | -0.103 | 0.102 |
| | Career satisfaction | | | | | 0.006(0.041) | -0.075 | 0.086 |
| | Community recognition | | | | | 0.047(0.04) | -0.126 | 0.033 |
| Work-family conflict | Research satisfaction | 0.187 | 0.003 | -0.132 | 0.000 | -0.025(0.009) | -0.045 | -0.008 |
| | Career satisfaction | | | -0.075 | 0.000 | -0.014(0.006) | -0.028 | -0.004 |
| | Community recognition | | | -0.071 | 0.001 | -0.013(0.006) | -0.026 | -0.003 |
| General support | Research satisfaction | 0.116 | 0.076 | 0.120 | 0.000 | 0.014(0.008) | -0.000 | 0.032 |
| | Career satisfaction | | | 0.124 | 0.000 | 0.014(0.008) | -0.000 | 0.032 |
| | Community recognition | | | 0.068 | 0.001 | 0.008(0.005) | -0.000 | 0.020 |
| Professional support | Research satisfaction | 0.059 | 0.314 | 0.024 | 0.381 | 0.001(0.002) | -0.002 | 0.007 |
| | Career satisfaction | | | 0.061 | 0.004 | 0.004(0.004) | -0.003 | 0.012 |
| | Community recognition | | | 0.025 | 0.244 | 0.002(0.002) | -0.002 | 0.007 |
| Financial support | Research satisfaction | 0.073 | 0.219 | 0.035 | 0.184 | 0.003(0.003) | -0.002 | 0.010 |
| | Career satisfaction | | | 0.001 | 0.947 | 0(0.002) | -0.004 | 0.004 |
| | Community recognition | | | 0.003 | 0.897 | 0(0.002) | -0.004 | 0.004 |
| Parents | | | | | | | | |
| (Direct effect) | Research satisfaction | | | | | -0.050(0.026) | -0.102 | 0.001 |
| | Career satisfaction | | | | | 0.054(0.023) | 0.009 | 0.100 |
| | Community recognition | | | | | -0.044(0.02) | -0.084 | -0.004 |
| Work-family conflict | Research satisfaction | 0.303 | 0 | -0.164 | 0.000 | -0.050(0.006) | -0.062 | -0.038 |
| | Career satisfaction | | | -0.146 | 0.000 | -0.044(0.006) | -0.056 | -0.034 |
| | Community recognition | | | -0.077 | 0.000 | -0.023(0.004) | -0.031 | -0.016 |
| General support | Research satisfaction | -0.247 | 0 | 0.113 | 0.000 | -0.028(0.005) | -0.037 | -0.019 |
| | Career satisfaction | | | 0.139 | 0.000 | -0.034(0.005) | -0.045 | -0.025 |
| | Community recognition | | | 0.077 | 0.000 | -0.019(0.003) | -0.026 | -0.013 |
| Professional support | Research satisfaction | 0.230 | 0 | 0.041 | 0.007 | 0.009(0.003) | 0.003 | 0.016 |
| | Career satisfaction | | | 0.024 | 0.075 | 0.006(0.003) | 0.000 | 0.011 |
| | Community recognition | | | 0.028 | 0.014 | 0.006(0.003) | 0.001 | 0.012 |
| Financial support | Research satisfaction | 0.295 | 0 | 0.003 | 0.768 | 0.001(0.004) | -0.006 | 0.008 |
| | Career satisfaction | | | -0.007 | 0.481 | -0.002(0.003) | -0.008 | 0.005 |

| Mediator | Outcome | a (gender → mediator) | p-value | b (mediator → outcome) | p-value | ab (path effect) or c' (direct effect, gender → outcome), Bootstrap standard error | 95% Percentile CI | |
|---|---|---|---|---|---|---|---|---|
| | | | | | | | Lower bound | Upper bound |
| | Community recognition | | | -0.002 | 0.846 | -0.001(0.003) | -0.006 | 0.006 |
| Objective career success | | | | | | | | |
| Non-parents | | | | | | | | |
| (Direct effect) | ARP | | | | | -0.003(0.09) | -0.179 | 0.174 |
| | ARC | | | | | -0.39(0.194) | -0.771 | -0.010 |
| | ARCo | | | | | 0.017(0.07) | -0.119 | 0.153 |
| Work-family conflict | ARP | 0.207 | 0.001 | -0.066 | 0.176 | -0.014(0.011) | -0.038 | 0.007 |
| | ARC | | | -0.150 | 0.066 | -0.031(0.019) | -0.074 | 0.000 |
| | ARCo | | | 0.028 | 0.445 | 0.006(0.008) | -0.008 | 0.024 |
| General support | ARP | 0.107 | 0.114 | 0.033 | 0.406 | 0.003(0.006) | -0.008 | 0.018 |
| | ARC | | | 0.205 | 0.037 | 0.022(0.017) | -0.002 | 0.062 |
| | ARCo | | | 0.048 | 0.133 | 0.005(0.005) | -0.002 | 0.017 |
| Professional support | ARP | 0.052 | 0.401 | -0.108 | 0.043 | -0.006(0.007) | -0.022 | 0.007 |
| | ARC | | | -0.074 | 0.427 | -0.004(0.009) | -0.026 | 0.010 |
| | ARCo | | | -0.080 | 0.057 | -0.004(0.005) | -0.016 | 0.005 |
| Financial support | ARP | 0.066 | 0.297 | -0.017 | 0.675 | -0.001(0.004) | -0.011 | 0.007 |
| | ARC | | | -0.191 | 0.004 | -0.013(0.013) | -0.042 | 0.010 |
| | ARCo | | | 0.046 | 0.270 | 0.003(0.004) | -0.004 | 0.014 |
| Parents | | | | | | | | |
| (Direct effect) | ARP | | | | | -0.335(0.077) | -0.486 | -0.185 |
| | ARC | | | | | -0.103(0.09) | -0.280 | 0.074 |
| | ARCo | | | | | 0.100(0.048) | -0.194 | -0.006 |
| Work-family conflict | ARP | 0.312 | 0 | 0.026 | 0.526 | 0.008(0.013) | -0.017 | 0.036 |
| | ARC | | | -0.017 | 0.750 | -0.005(0.018) | -0.043 | 0.028 |
| | ARCo | | | 0.019 | 0.323 | 0.006(0.007) | -0.007 | 0.020 |
| General support | ARP | -0.220 | 0 | 0.078 | 0.025 | -0.017(0.007) | -0.031 | -0.004 |
| | ARC | | | 0.071 | 0.174 | -0.016(0.011) | -0.037 | 0.004 |
| | ARCo | | | 0.044 | 0.030 | -0.010(0.004) | -0.018 | -0.002 |
| Professional support | ARP | 0.234 | 0 | -0.054 | 0.146 | -0.013(0.009) | -0.031 | 0.004 |
| | ARC | | | 0.015 | 0.787 | 0.004(0.013) | -0.023 | 0.030 |
| | ARCo | | | -0.034 | 0.092 | -0.008(0.005) | -0.018 | 0.002 |
| Financial support | ARP | 0.278 | 0 | -0.006 | 0.862 | -0.002(0.011) | -0.023 | 0.020 |
| | ARC | | | -0.028 | 0.572 | -0.008(0.013) | -0.035 | 0.017 |
| | ARCo | | | -0.001 | 0.956 | 0(0.005) | -0.011 | 0.010 |

#### Survey Questions

Note: Only questions supporting analyses in this study are listed below.

With which gender do you most identify?
- Female
- Male
- Other, please specify
- Prefer not to disclose

Please specify your ethnicity:
- White
- Black or African American
- American Indian or Native American
- Asian or Pacific Islander
- Hispanic or Latino
- Other, please specify
- Prefer not to disclose

What is your current marital status? This includes marriage or domestic partnership with the opposite or same sex. In this context, a domestic partnership includes those who are officially registered or who have lived together for more than 2 years.
- Never married or never taken part in a domestic partnership
- Never married but took part in a domestic partnership
- Married, or in a domestic partnership
- Separated
- Divorced
- Widowed
- Not list, please specify

What is/are your general area(s) of study or research? Check all that apply.
- Arts & Humanities
- Medical Sciences
- Natural Science & Engineering
- Social Sciences
- Other, please specify

What is your current rank or role? Check all that apply. - Student (Bachelor, Master, or Doctoral)
- Student (Bachelor, Master, or Doctoral)
- Post-doctoral fellow
- Lecturer (teaching graduate or undergraduate courses)

- Technician or technician assistant (e.g., statistician, laboratory assistant)
- Research associate (at a public or private institution)
- Senior researcher (at a public or private institution)
- Assistant professor
- Associate professor
- Full professor
- Emeritus professor
- Other, please specify

This project focuses on the relationship between the career development and familial role of researchers. If you are currently a student and never had any work experience, you have the option to quit the survey.

- I would like to quit the survey.
- I would like to continue the survey

How many children (of all ages) do you have, including step-, adopted, and biological children?

- 0
- 1
- 2
- 3
- 4
- 5
- 6 or more

Is the number of children (include 0) you currently have related to your career considerations (more or less)?

- Yes
- No
- Prefer not to disclose

**(For those who have children)** Please evaluate the overall impact of child-rearing on your career development:

- Negative
- Slightly negative
- Almost no influence
- Slightly positive
- Positive

**(For those who have ever been married or cohabited)** Does/Did your current or most recent spouse/partner's primary job duties include conducting research?

- Yes
- No

- I am not sure
- Not applicable

**(For those who answered Yes in the previous question)** Did you ever collaborate on research projects with your current or most recent spouse/partner?

- Yes
- No
- Not applicable

**(For those who have ever been married or cohabited)** Overall, to what extent did your current or most recent spouse/partner provide the following support to your career development?

| | Not at all | A little bit | Moderate | Substantial |
|---|---|---|---|---|
| Financial support (such as providing reasonable financial support when needed) | ○ | ○ | ○ | ○ |
| Emotional support (such as listening to your complaints, giving you a pep talk when needed) | ○ | ○ | ○ | ○ |
| Time support (such as helping you take care of children and letting you focus more on work) | ○ | ○ | ○ | ○ |
| Decision support (such as your spouse/partner agreeing to move when you are offered an opportunity to do so for a job in another city) | ○ | ○ | ○ | ○ |
| Technical support (such as helping you solve problems in your work, or participating in your research) | ○ | ○ | ○ | ○ |
| Network support (such as introducing people to you who might benefit your research) | ○ | ○ | ○ | ○ |
| Other support, please specify | ○ | ○ | ○ | ○ |

**(For those who have ever been married or cohabited)** Have you experienced any of the followings that impeded your career development because of spouse/partner related reasons?

| | Not at all | A little bit | Moderate | Substantial |
|---|---|---|---|---|
| Emotional pressure (e.g., does not care about your career; makes you feel more depressed when you have problems at work) | ○ | ○ | ○ | ○ |
| Career-family conflict (e.g., you sacrificed your own working | ○ | ○ | ○ | ○ |

| | | | | |
|---|---|---|---|---|
| time/career opportunity to take care of family) | | | | |
| Decision nonsupport (e.g., does not agree to you accepting a more promising job in another city) | ○ | ○ | ○ | ○ |
| Marital dissatisfaction (e.g., you couldn't concentrate on work due to unresolved marital conflict) | ○ | ○ | ○ | ○ |
| Network constraints (e.g., limits your involvement with the opposite sex which may benefit your career development) | ○ | ○ | ○ | ○ |
| Other, please specify | ○ | ○ | ○ | ○ |

**(For those who have ever been married or cohabited)** Please rate the family-work conflicts you have experienced, if any.

| | Not at all | A little bit | Moderate | Substantial |
|---|---|---|---|---|
| Time-based conflict (e.g. working late on weekends to complete a work project, interfering with the time you can spend with family) | ○ | ○ | ○ | ○ |
| Strain-based conflict (e.g. an employee is not able to concentrate on work because he/she is concerned about his/her sick child/spouse/partner) | ○ | ○ | ○ | ○ |
| Behavior-based conflict (e.g. high-level employees are expected to be aggressive and unyielding at work but kind and considerable with his/her spouse/children) | ○ | ○ | ○ | ○ |

I am satisfied with the progress I have made towards meeting my research achievement goals.

- Strongly disagree
- Disagree
- Somewhat disagree
- Neither agree nor disagree
- Somewhat agree
- Agree
- Strongly agree
- Not applicable

I am satisfied with the progress I have made towards meeting my career achievement goals.

- Strongly disagree
- Disagree

- Somewhat disagree
- Neither agree nor disagree
- Somewhat agree
- Agree
- Strongly agree
- Not applicable

I have been recognized for my contributions to scholarly communities.

- Strongly disagree
- Disagree
- Somewhat disagree
- Neither agree nor disagree
- Somewhat agree
- Agree
- Strongly agree
- Not applicable
